# Consequence of adjustments for demographic or clinical covariates and a recommended solution in genome-wide association studies

**DOI:** 10.1101/2021.12.07.471675

**Authors:** Xiaoru Sun, Hongkai Li, Yuanyuan Yu, Zhongshang Yuan, Chuandi Jin, Lei Hou, Xinhui Liu, Qing Wang, Fuzhong Xue

**Author notes:** Correspondence to: Fuzhong Xue. These authors contributed equally to this work (Xiaoru Sun, Hongkai Li). Postal Address: Institute for Medical Dataology, Shandong University, No.44 Wenhua West Road, Jinan, Shandong, 250012, China.

## Abstract

Genome-wide association study (GWAS) is fundamentally designed to detect disease-causing genes. To reduce spurious associations or improve statistical power, about 80% of GWASs arbitrarily adjusted for demographic or clinical covariates. However, adjustment strategies in GWASs have not achieved consistent conclusions. Given the initial aim of GWAS that is to identify the causal association between a specific causal single-nucleotide polymorphism (SNP) and disease trait, we summarized all complex relationships of the target SNP, covariate and disease trait into 15 causal diagrams according to various roles of the covariate. Following each causal diagram, we conducted a series of theoretical justifications and statistical simulations. Our results demonstrate that it is unadvisable to adjust for any demographic or clinical covariates. We illustrate our point by applying GWASs for body mass index (BMI) and breast cancer, including adjusting and non-adjusting for age and smoking status. Genetic effects and P values might vary across different strategies. Instead, adjustments for SNPs (*G*′) should be strongly recommended when *G*′ are in linkage disequilibrium with the target SNP, and correlated with disease trait conditional on the target SNP. Specifically, adjustment for such *G*′ can block all the confounding paths between the target SNP and disease trait, and avoid over-adjusting for colliders or intermediaries.

## Introduction

Genome-wide association study (GWAS) is fundamentally designed to detect disease-causing genes since it was proposed in 2005 (Klein 2005), which is aimed to confirm the causal associations between single-nucleotide polymorphisms (SNPs) and disease traits (Hu et al. 2018; Visscher et al. 2012). Theoretically, the causal direction is natural from SNP to disease trait (*Y*), and none of demographic or clinical covariates (*C*) alter individual’s born genotype (i.e. SNP). In addition, although environmental factors or the cumulative number of stem cell divisions could lead to genomic mutation during one’s lifespan at the single-cell level (Tomasetti et al. 2017; Tomasetti and Vogelstein 2015), its genotyping data in GWAS is actually averaged from plenty of cells with extremely infrequent mutation for a specific SNP which is still the born one. Therefore, the confounding path SNP ← *C* → *Y* is not intuitively formed, and causal association analysis between the target SNP and disease trait is unbiased without adjusting for *C*. In this situation, univariate regression models and Cochran–Armitage trend test were widely used to identify causal SNPs at the early GWASs (Franke et al. 2010). Subsequently, in order to control the confounding bias due to population stratification (Campbell et al. 2005; Freedman et al. 2004; Helgason et al. 2005; Hirschhorn and Daly 2005; Lander and Schork 1994; Lohmueller et al. 2003; Marchini et al. 2004; Thomas et al. 2005), a series of methods such as genomic control (Devlin et al. 2004; Devlin and Roeder 1999), genomic principal components regression (Price et al. 2006) were essentially added to the univariate models. In this way, GWAS has successfully identified some novel variant–trait associations and discovered its biological mechanisms, and these findings have diverse clinical applications. Also, GWAS has provided insight into the ethnic variation of complex traits. Recently, low-frequency and rare variants were detected with the developments of next-generation sequencing technology by GWAS strategies, and numbers of genetic variants including monogenic or oligogenic disease genes, copy number variants are expected to identify. These undoubtedly rise our confidence in post-GWAS, and GWAS data is used for multiple applications beyond gene identification (Tam et al. 2019).

Nevertheless, in the published GWASs, various demographic or clinical covariates were arbitrarily adjusted for using multiple regression models, claiming to reduce spurious associations or improve statistical power (Aschard et al. 2015; Bush and Moore 2012). This adjustment strategy has been controverted by statistical geneticists (Aschard et al. 2015; Pirinen et al. 2012; Schisterman et al. 2009; Zhang et al. 2018). Although these adjustments could improve statistical power to some extent under linear regression model for quantitative traits, it is easy to make mistakes by adjusting for colliders (Day et al. 2016) or intermediaries (Aschard et al. 2015) unknowingly. These mistaken adjustments usually lead to underestimate the effects of SNPs on traits, which are less than helpful for discovering causal SNPs. Meanwhile, some studies found the adjustments would reduce statistical power rather than improving it for qualitative traits under logistic regression model (Pirinen et al. 2012). Unfortunately, these arbitrary adjustments have not raised enough attentions, even it has become a common view that demographic or clinical covariates have to be adjusted for in GWASs.

Considering the fundamental objective of GWASs is to identify causal SNPs on disease traits, we believe the priority should not consider improving the statistical power before we obtain the unbiased effects of causal SNPs on disease traits. Meanwhile, false discovery rate (FDR) should be simultaneously controlled in order to avoid the dilemma that the higher the power is, the higher the FDR is. This will inevitably lead to large obstacles in the subsequent experimental validations. Although the roles of the target SNP (*G*), covariate (*C*) and disease trait (*Y*) as well as their relationships are extremely complex, given the initial aim of GWASs, we simplified and summarized all the scenarios into 15 causal diagrams according to the various roles of *C* (Fig. 2). Furthermore, in order to compare the performances of adjustment and non-adjustment for *C*, combining graphical model (Pearl 2000) with counterfactual model (Petersen and van der Laan 2014), we conducted a series of theoretical justifications and simulations under linear model and logistic model following each causal diagram in Fig. 2, respectively. We illustrate our point by applying GWASs for body mass index (BMI) and breast cancer, including adjusting and non-adjusting for age and smoking status. Genetic effects and P values might vary across different strategies. We explore this phenomenon in detail and discuss the ramifications for future genome-wide association studies of correlated traits and diseases.

## Results

### The landmarks of adjustments for covariates in GWASs

We summarized 1686 ones relevant to 12 types of traits from 3198 published GWASs papers from March 2005 to July 2018 on the GWAS Catalog to highlight the landmarks of adjustments for covariates (Fig. 1). Obviously, adjustments for demographic or clinical covariates have been becoming the mainstream since 2007, which accounted for 79.2% of the whole GWAS papers (Fig. 1a). Among these adjustment methods, multiple linear regression models and multiple logistic regression models were widely used with a proportion of 25.4% and 37.6%, respectively (Fig. 1b). In exception for necessary adjustments for regions and principal components to correct for population stratification, dozens of demographic or clinical covariates were adjusted. The most commonly adjusted covariates were age and gender, followed by body mass index, smoking status, drinking status, diabetes, hypertension and tumor, etc. Specifically, SNPs correlated to the target SNP were also adjusted for in several papers (Tintle et al. 2015; Leng et al. 2015) (Fig. 1c). Under the above analytic strategies, only 2.83% of the total identified 23,938 SNPs in the 1686 published GWAS papers were successfully validated by experiments (Fig. 1d). Such a definitely low proportion of verification motivates us to have to rethink the rationality of the current analytic strategies in GWASs, especially for the arbitrary adjustments for demographic or clinical covariates. Theoretical analysis

**Fig. 1.**
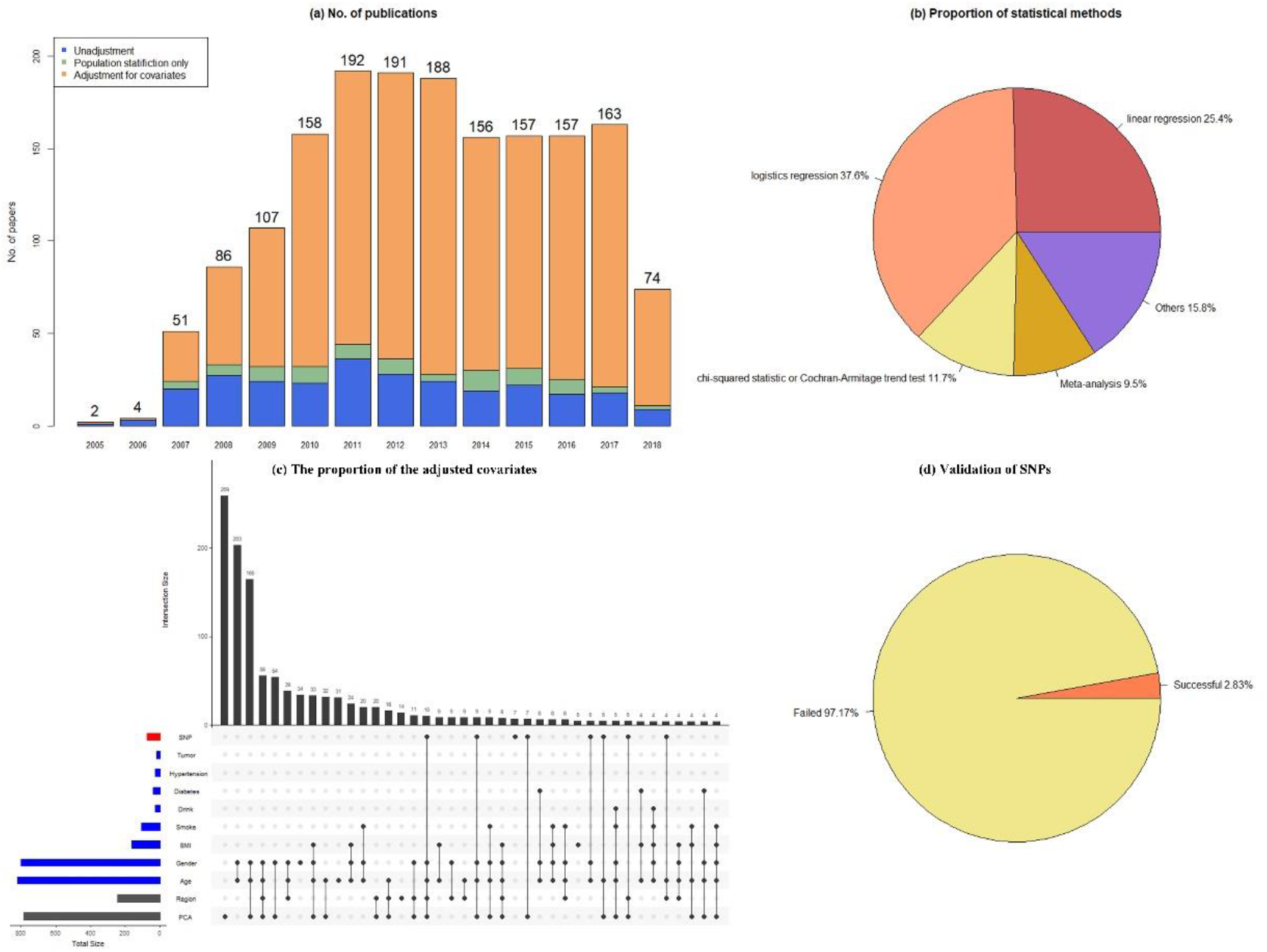
The statistical summary of the adjustment for covariates. (a) The number of published papers from March 2005 to July 2018. (b) The proportion of commonly used statistical methods in GWASs. (c) The 11 most commonly adjusted covariates in published GWASs. (d) The proportion of validated SNPs in the published GWASs.

We conducted a series of theoretical analyses under the linear model and logistic model based on the causal diagrams in Fig. 2, respectively. A linear model is applied to estimate the effect of *G* on quantitative trait *Y*. For simplicity, we assume *G* and *C* have been standardized with mean zero and variance one (hence, the covariance of *G* and *Y*, the correlation coefficient and the regression coefficient are equal, i.e. *σ*_*YG*_ = *ρ*_*YG*_ = *β*_*YG*_). Then regression coefficient *β*_*YG*_ and partial regression coefficient *β*_*YG*·*C*_ can be calculated by

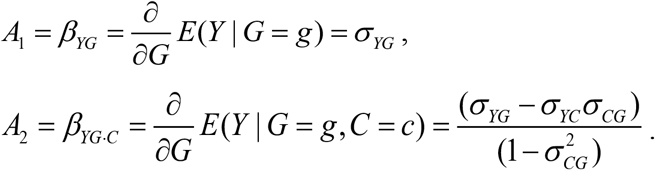

The true causal effect 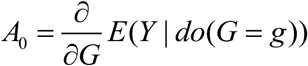 is calculated based on *do calculus* proposed by Pearl (Pearl 2000). Then the bias could be denoted by *bias*_1_ = *A*_1_ − *A*_0_ when *C* is not adjusted for, while *bias*_2_ = *A*_2_ − *A*_0_ is the bias with *C* adjusted. Details are shown in the Supplementary materials. The theoretical results can be summarized as the following theorems.

#### Theorem 1.

The marginal effect and conditional effect are both equal to the causal effect of the SNP *G* on the trait *Y* under linear model before and after adjusting for *C*, if

1. *G* ⊥ *C* and 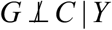; or
2. 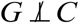 and *G* ⊥ *C* | *G*

*Proof*. The true causal effect of *G* on *Y* can be calculated as 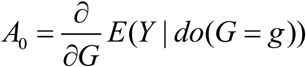 according to the *do-calculus* under linear model. If the *G* ⊥ *C* and 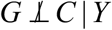, or 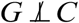 and *Y* ⊥ *C* | *G*, C is not on the back-door path of G→Y. Then 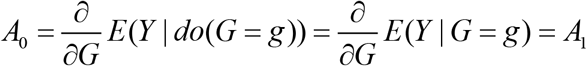.

The conditional effect 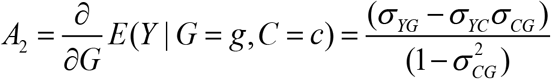. If *G* ⊥ *C, σCG* = 0 *A*_2_ = *σ*_*YG*_ = *A*_1_. If 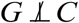 and *Y* ⊥ *C* | *G, σ*_*YC*_ = *σ*_*YG*_*σ* _*CG*_, *A*_2_ = *σ*_*YG*_ = *A*_1_. Thus the marginal effect and conditional effect are both equal to the causal effect of the *G* on *Y*.

For the qualitative trait Y, the logistic model is applied for causal effect estimation. The theoretical results can be summarized as followings. And the proof is shown in the supplementary materials.

#### Theorem 2.

The marginal effect and conditional effect are both equal to the causal effect of the SNP G on the trait Y under logistic model before and after adjusting for C, if

1. *Y* ⊥ *C* | *Y*; or
2. *Y* ⊥ *C* | *G*

If *C* is a non-confounding factor and the above conditions are not satisfied, a new bias will be induced with adjusting for *C*. If C is confirmed to be a confounder, adjusting for C can eliminate confounding bias under the linear model and decrease bias under the logistic model.

### Simulations: the consequences of adjustments for covariates in GWASs

We conducted a series of simulations under linear model and logistic model following each causal diagram in Fig. 2, respectively. Three aspects of evaluation metrics, including bias and standard error (SE) for assessing the accuracy and precision, Type I error and power for stability and statistical efficiency of model, FDR, true discovery rate (TDR) and Matthews correlation coefficient (MCC) for performances of causal SNP’s identification, were simultaneously used to evaluate the results before and after adjusting for *C* with various roles (Supplementary Notes).

**Fig. 2.**
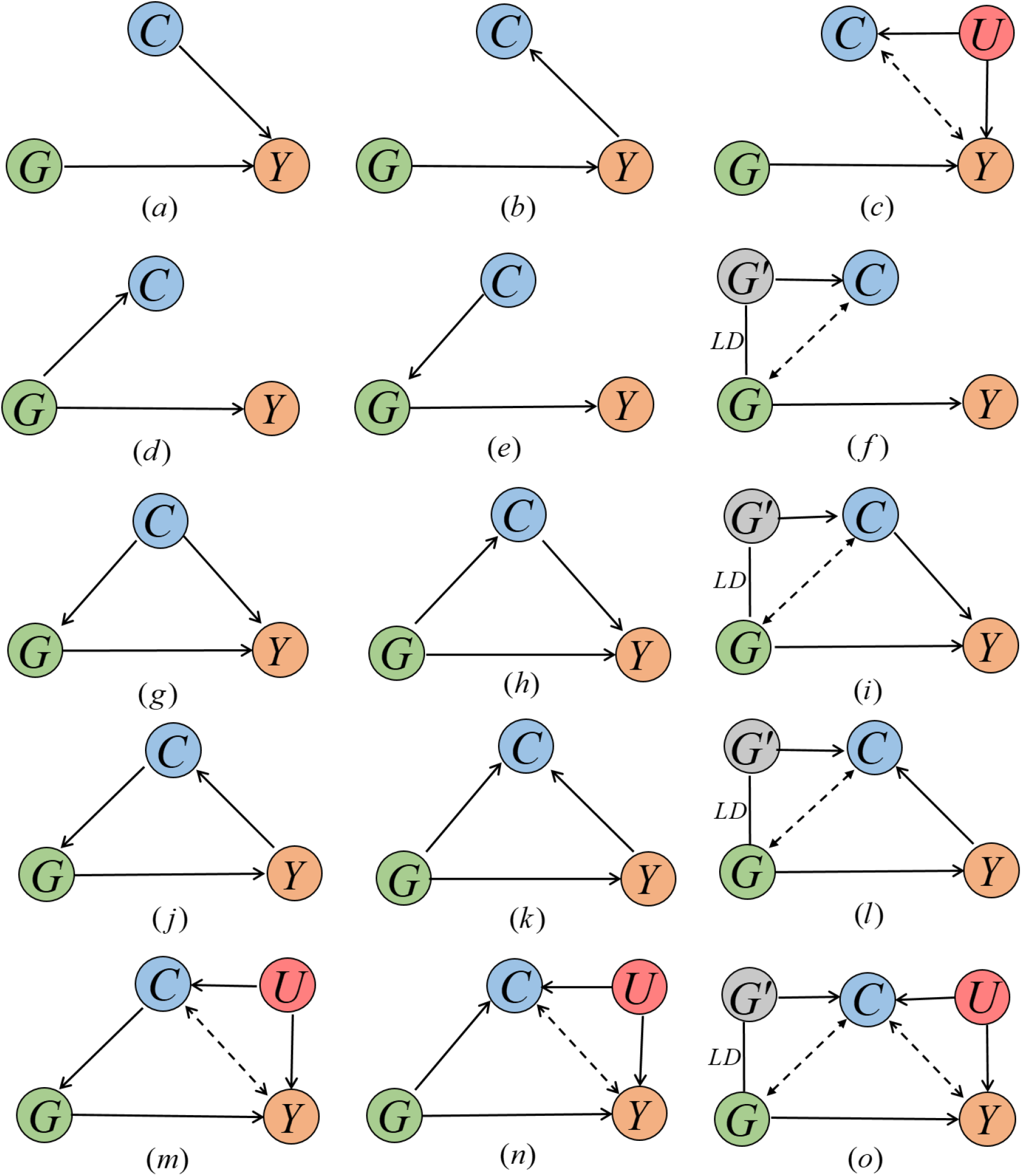
15 causal diagrams presenting all the complex relationships of the target SNP (*G*), covariate (*C*) and disease trait (*Y*) given the initial aim of GWAS. *G* and *Y* are the interested target SNP and disease trait, respectively. (***a-c***) *C* has no direct causal association with *G*. (***d-f***) *C* has no direct causal association with *Y*. (***g-i***) *C* is a cause of *Y* and correlated with *G*. (***j-l***) *C* is caused by *Y* and correlated with *G*. (***m-o***) *C* is correlated with *Y* via the unobserved variable *U*.

#### C has no direct causal association with G (Fig. 2 a-c)

For Fig. 2*a*, theoretically, we both obtained unbiased effects of *G* on *Y* before and after adjusting for *C* under linear model (Supp. S1), while simulations indicated that precision and statistical power improved after adjusting for *C* (Supp. S2). Unfortunately, as an absolutely common pattern in genomics, when the target SNP and other SNPs were in high linkage disequilibrium (LD), higher FDR was also observed (usually more than 95%). As a consequence, MCC reduced and gained little benefit from adjustment for *C*. Even the target SNP and other SNPs were in low LD, the distinction of two MCCs between adjustment and non-adjustment for *C* became less significant as the effect of *G* on *Y* got larger, and thus adjustment for *C* obtained little benefit (Supp. S3). On the other hand, when adjusting for *C* under logistic model, bias occurred due to collapsibility (Guo and Geng 1995) (Supp. S4). In this situation, adjustment for *C* decreased precision, little improved power, expanded FDR, reduced TDR and finally lowered MCC in comparison with non-adjustment (Supp. S5-6). In summary, adjustment for *C* should not be encouraged. For Fig. 2*b*, an uni-path colliding bias of the effect from *G* to *Y* was induced after adjusting for *C* under linear model in that *G* and *Y* collide at *C* (Supp. S7). Simulation results revealed that a slightly greater precision but larger FDR, lower power, TDR and MCC were obtained with adjusting for *C* (Supp. S8-S9). Although unbiased effects were obtained before and after adjusting for *C* under logistic model, the precision reduced when adjusting for *C* (Supp. S10-S11), and same consequences of other metrics occurred as those in linear model (Supp. S12). Unbiased effect of *G* on *Y* was obtained with adjusting for *C* under linear model but a small bias under logistic model when the effect of *G* on *Y* became large in Fig. 2*c* (Supp. S13, S16). In this scenario, even though precision and power improved, MCC benefited little (Supp. S14-S15, S17-S18). In summary, adjustment for *C* is not recommended, if no direct causal association exists between *C* and *G*.

#### C has no direct causal association with Y (Fig. 2 d-f)

Since Fig. 2*e* does not exist actually, we just take Fig. 2*d* and 2*f* into consideration here. Both adjustment and non-adjustment for *C* obtained unbiased effects of *G* on *Y* under linear and logistic models in Fig. 2*d* and Fig. 2*f* (Supp. S19, S22, S25 and S28). While all the other evaluation metrics of adjustment for *C* were lower than or equal to that without adjustment (Supp. S20-S21, S23-S24, S26-S27 and S29-S30), thus adjustment for *C* is not reasonable.

#### C is a cause of Y and correlated with G (Fig. 2 g-i)

As *C* does not affect *G* theoretically, Fig. 2*g* is not under consideration. Obviously, adjustment for intermediate variable *C* in Fig.2*h* led to a bias both under linear and logistic models (Supp. S31, S34). Due to the biased estimation together with the collinearity between *G* and *C*, all the evaluation metrics went worse (Supp. S32-S33, S35-S36). In theory, adjustment for confounder *C* or *G*′ in Fig. 2*i* under linear model obtained unbiased effect of *G* on *Y* (Supp. S37). Both FDR and TDR decreased, then MCC increased though the statistical power decreased (Supp. S38-S39). However, it should be noted that we are usually hard to distinguish the roles of *C* from confounder, intermediary and collider in practice. Hence we strongly suggest adjust for *G*′ rather than *C*. In spite of biased effects of *G* on *Y* before and after adjusting for *C* or *G*′ under logistic model in presence of collapsibility (Supp. S40), adjustment for *G*′ performed better than non-adjustment and adjustment for *C* (Supp. S41-S42).

#### C is caused by Y and correlated with G (Fig. 2 j-l)

As the hypotheses that *C* is not a cause of *G* makes no sense, Fig. 2*j* is not considered here. Adjustment for the collider *C* in both Fig. 2*k* and Fig. 2*l* necessarily led to underestimated effects of *G* on *Y* under linear and logistic models (Supp. S43-S44, S46-S47, S49-S50, S52-S53). As a result, all the three aspects of evaluation metrics went much worse (Supp. S43-S54).

#### C is correlated with Y via the unobserved U (Fig. 2 m-o)

For Fig. 2*m*, there is no need to take into account. For the other two scenarios Fig. 2*n* and Fig. 2*o*, since *C* still works as a collider, adjustment for it incurred the same consequences as those in Fig. 2*k* and Fig. 2*l* (Supp. S55-S66).

## Applications

### Influence of adjusted covariates on GWAS results of BMI

BMI information was collected from 360,108 participants after excluding samples with missing height and weight information. SNPs with MAF < 0.01, call rate < 0.95 or P < 1.0×10 ^-10^ from a test for Hardy-Weinberg equilibrium were also excluded. In view of population stratification bias, the first 10 principal components of genetic information were adjusted to correct the stratification bias during genome-wide association analysis. 48982 SNPs were significant associated with BMI (P < 5.0×10 ^-8^) without adjusting for age and smoking. Fig. 3a showed the comparison of the P values before and after adjusting for age and smoking. A few SNPs got a smaller P values after adjustment for covariates. But the overall impact is small.

**Fig. 3.**
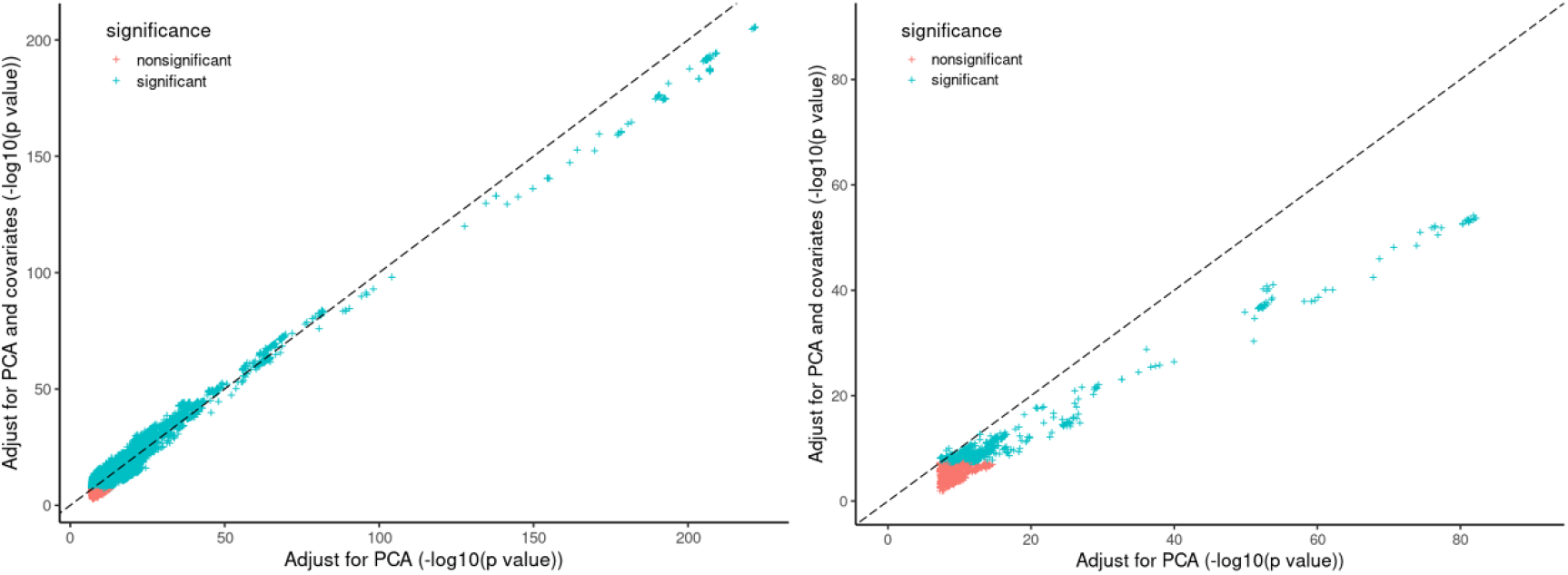
The comparison of the P values before and after adjusting for covariates. (a) GWAS results of BMI. (b) GWAS results of breast cancer.

### Influence of adjusted covariates on GWAS results of breast cancer

We collected 13,192 patients with breast cancer and 153,306 healthy participants for GWAS. The first 10 principal components of genetic information were adjusted. 2074 SNPs were significant associated with breast cancer (P < 5.0×10 ^-8^) without adjusting for age and smoking. After adjusting for age and smoking, only 1020 SNPs were still significant. Most P values were increased after the adjustment, which indicates the loss of precision of models (Fig. 3b).

## Discussions

GWAS is one of the most important tools for discovering susceptibility loci of traits and diseases. The rationality of adjustment for covariates in GWAS has been discussed for a long time (Aschard et al. 2015; Pirinen et al. 2012; Schisterman et al. 2009; Zhang et al. 2018). To make this problem clear, we conducted a series of theoretical justifications and simulations to clarify the consequences of adjustment for covariates in GWAS. We found no rationality for adjusting for any demographic or clinical covariates.

For the 15 causal diagrams in Fig. 2, the covariate *C* is a non-confounding factor except in Fig. 2i. In Fig. 2a, *C* is an independent cause of *Y*. Adjusting for *C* in this scenario can improve the precision without biasing the effect estimation under linear model^30^. That is the reason for the proposal of adjustment in GWAS. However, high precision may not benefit to GWAS because of the accompanying high FDR. Few research discusses this problem under logistic model, which is the most frequently used model for qualitative trait. Geng et al. discussed the collapsibility of logistic model (Guo and Geng 1995). But the biased estimation and decreased precision are not discussed under logistic model for this scenario. If *C* is independent of *G* and not causally associated with *Y* (Fig. 1c), adjustment has a similar performance to Fig. 2a.

In Fig. 2(b, k, l, n, o), *C* is a collider of *G* → *Y*. Adjustment for *C* is not advisable. For the uni-path in Fig. 2b, *C* is a special collider. Adjustment for *C* can obtain an unbiased OR estimation due to the collapsibility of logistic model. But the loss of the precision is not unavoidable. Thus adjustment for *C* is not recommended.

*C* in Fig. 2d and Fig. 2f are only directed correlated with *G*. It is labelled as unnecessary adjustment in these two scenarios under linear model^23^. In this study, we complemented the performance of adjustment under logistic model. It is not recommended for adjustment due to the loss of precision. In Fig. 2h, *C* is a mediation. Overadjustment for *C* would miss the indirect effects of *G* on *Y*.

The confounding path of *G* → *Y* in Fig. 2i is induced by the correlation of G and *G*′, *G*′ and *C. G*′ is an equivalent confounder of *C* in this scenario. Adjustment for *G*′ can obtain the same unbiased estimation as the performance of *C*. Confounding *C* is hard to be distinguished from the collider and mediation. While the confounding *G*′ can be selected according to the associations with *G, C* and *Y*.

Based on our theoretical justifications and statistical simulations as above, we conclude that it is not sensible to adjust for any demographic or clinical covariates, instead adjustments for SNPs (*G*′) should be strongly recommended when the following requirements that *G*′ are in linkage disequilibrium with the target SNP, and correlated with disease trait conditional on the target SNP, are satisfied. Specifically, adjustment for *G*′ can block all the confounding paths (*G − G*′ … *C* → *Y* and *G −G*′ → *Y*) between the target SNP and disease trait from the perspective of the causal inference (Fig. 4). Furthermore, such strategy can simultaneously avoid mistakenly adjusting for collider or intermediaries unknowingly. Subsequently, further research should mainly focus on how to adjust for *G*′.

**Fig. 4.**
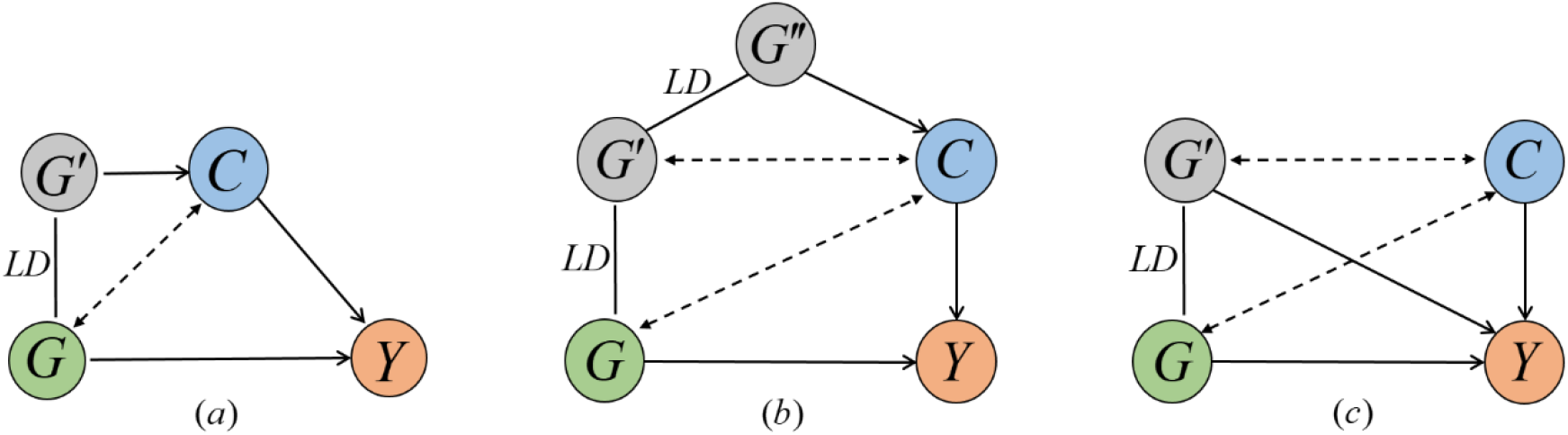
Causal diagrams of three scenarios that *G*′ is recommended to be adjusted for. *G* and *Y* are the interested target SNP and disease trait, respectively. *G*′ and *G* are in LD. *C* is a cause of *Y* and correlated with *G*. (*a*) is the same scenario as Fig. 2*i*. *G − G*′ → *C* → *Y* is a confounding paths from *G* to *Y*. (*b*) is a more complex scenario that *G − G*′ *− G*′′ → *C* → *Y* is the confounding paths from *G* to *Y. G*′ is in LD with *G*′′ and correlated with *C*. (*c*) is the scenario that *G − G*′ *− C* → *Y* and *G − G*′ → *Y* are two confounding paths from *G* to *Y. G* and *G*′ are correlated with *C*.

## Methods

### Search strategy and inclusion criteria of published GWAS papers

We searched GWAS Catalog (https://www.ebi.ac.uk/gwas/) for the published GWAS papers from March 2005 to July 2018. After excluding duplications papers, there are 3198 published GWASs papers. Among them, 1686 papers are included, which involve in 12 representative fields: biological process, cardiovascular measurement, digestive system disease, immune system diseases, hematological measurement, lipid measurement, lipoprotein measurement, nervous system disease, metabolic diseases, inflammatory marker measurement, liver enzyme measurement and response to drug. Theoretical justification for the biases of adjustment for *C*

Biases of effects before and after adjusting for *C* under linear model. A linear model is applied to obtain the effect of *G* on the quantitative trait *Y*. In order to determine if we obtain unbiased effect adjustment or non-adjustment for *C*, the following quantities are compared: 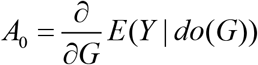 is the incremental causal effect of *G* on *Y* according to the *do calculus* (Pearl 2000); 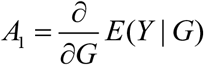 is the incremental unconditional dependence of *G* on *Y*, given by the regression coefficient of *G* on *Y*; 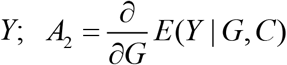 is the incremental conditional dependence of *G* on *Y*, given by the coefficient of *G* in the regression of *Y* on *G* and *C*. The unadjusted bias could be denoted by *bias*_1_ = *A*_1_ − *A*_0_, while *bias*_2_ = *A*_2_ − *A*_0_ is the adjusted bias.

Biases of effects before and after adjusting for *C* under logistic model. Assuming the qualitative trait *Y* is binary, logistic model is used to obtain the effect of *G* on *Y*. To infer if we obtain unbiased effects before and after adjusting for *C, bias*_1_ = *A*_1_ − *A*_0_ and *bias*_2_ = *A*_2_ − *A*_0_ are obtained through the following quantities

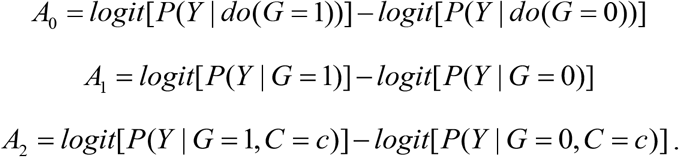

Simulations

Simulations are performed for bias, SE, Type I error and statistic power under linear model. Assuming *C* and *G* have been standardized with zero-mean and unit-variance. For instance, in Fig. 2a, *Y* is generated as *Y* = *β*_0_*G* + *β*_1_*C* + *ε*_1_, where *β*_0_ is the conditional effect of *G* on *Y, β*_1_ the conditional effect of *C* on *Y, ε*_1_ the random term, *ε*_1_ ∼ *N* (0,1).

For each causal diagram, performances of the two models: 1) non-adjustment for the extraneous factor *C*, 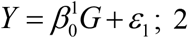) adjustment for *C*, 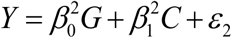 are compared. For each scenario, given other parameters remain constant (i.e. *β*_0_ = 0 and *β*_1_ = 0.1), we simulate to obtain type I error when the range of sample size changes from 500 to 5000 with 500 as a gap. Then bias 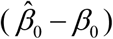, standard error 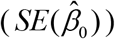 and statistical power are calculated as *β*_0_ = 0.1, *β*_1_ = 0.1 and sample size ranges from 500 to 5000. Finally, we set sample size at 2000 and *β*_1_ = 0.1, bias, standard error and statistical power are obtained as *β*_0_ changes from 0.05 to 0.9 with 0.05 as a gap. Similarly, the sample size is set at 2000 and *β*_0_ = 0.1, *β*_1_ changes from 0.05 to 0.9 with 0.05 as a gap. 1000 simulations are repeated for each scenario.

Simulations are performed for FDR, TDR and MCC under linear model. Following the LD patterns in the whole genome data, 22 genomic regions (each with 50 SNPs) are selected from each of 22 autosomes. Then a merged mimic genomic region with 1100 SNPs is created for our simulation. In this case, this simulated region is a miniature of whole genome, and covers the characteristics of the real world genome data. Software gs2.0 is used to generate the genotype data of 100,000 individuals referring to HapMap phase III CEU data (http://hapmap.ncbi.nlm.nih.gov/). The genotype data is coded by the additive genetic model.

Subsequently, two simulation projects are designed to assess the performances of causal SNP’s identification by three commonly-used metrics: false discovery rate (FDR), true discovery rate (TDR) and Matthews correlation coefficient (MCC).

In simulation project 1, 10 independent SNPs (*G*) causing the disease trait (*Y*) are generated following the Bernoulli distributions *P*(*G* = 0, *G* = 1, *G* = 2) = (*p*^2^, 2 *pq, q*^2^). Then the 10 causal SNPs are randomly inserted into 10 ones from 22 autosomes in the simulated regions, respectively. Given *G* based on each causal diagram, we generate the quantitative trait *Y* and 10 corresponding *C*. Linear model is used to calculate FDR, TDR and MCC after adjusting the *P* value of all the SNPs by Bonferroni correction.

In simulation project 2, each of 10 SNPs (*G*) causing *Y* are randomly selected from 10 simulated regions from different autosomes with high LD structure (most pair of SNPs with a correlation coefficient r^2^ > 0.6), respectively. The corresponding *C* and *Y* are generated based on each causal diagram. Performances of the two models adjustment and non-adjustment for *C* are evaluated as the same as project 1.

Simulations are performed for bias, SE, Type I error and statistic power under logistic model. Assuming *Y* is qualitative trait following the Bernoulli distributions. Taking Fig. 2a as an example again, we randomly generate *G, C* and *Y* with

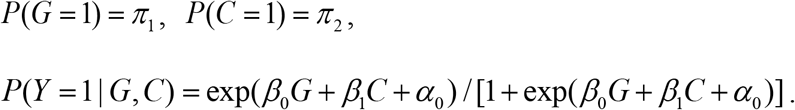

For each scenario, we also establish two models to compare their performances:

1. Non-adjustment for *C*, 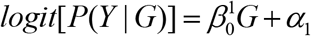;
2. Adjustment for *C*, 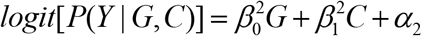.

The subsequent simulation procedures are the same as those under the linear model. The only difference is that the odds ratio (*OR*) between two variables starts from 1.1 to 2 with 0.1 as the gap, then 2.5, 3 and 4.

Simulations are performed for FDR, TDR and MCC under logistic model. We assume qualitative trait *Y* is a binary variable and logistic model is used. Using the above genomic data, FDR, TDR and MCC are calculated in the same way as the linear model.

All simulation studies are conducted using software R from CRAN (http://cran.r-project.org/).

## Applications

To illustrate our point, we applied GWAS analyses for body mass index (BMI) and breast cancer, including adjusting and non-adjusting for age and smoking status. Genetic effects and P values might vary across different strategies. We explore this phenomenon in detail and discuss the ramifications for future genome-wide association studies of correlated traits and diseases.

## Author contributions

F.X., X.S. and H.L. jointly conceived the idea and designed the study. X.S. conducted the literature review, the theoretical justifications and simulation studies, then finished the draft of the manuscript. Y.Y. and Z.Y. contributed to the design of the study. C.J., Q.Z., L.H. and X.L. reviewed and summarized GWAS papers. F.X., H.L. and Q.W. revised the manuscript. All authors read and approved the final manuscript.

## Competing interests

The authors declare no competing interests.

## Additional information

Details of the theoretical derivation and simulation results are shown in Supplementary Information.docx.

## Funding

This work was supported by the National Natural Science Foundation of China (81773547 and 82003557), National Key Research and Development Program of China (2020YFC2003500) and Shandong Provincial Natural Science Foundation of China (ZR2019ZD02).

## References

Aschard H, Vilhjálmsson BJ, Joshi AD, Price AL, Kraft P (2015) Adjusting for Heritable Covariates Can Bias Effect Estimates in Genome-Wide Association Studies. The American Journal of Human Genetics 96: 329–339

Bush WS, Moore JH (2012) Chapter 11: Genome-wide association studies. PLoS Comput Biol 8: e1002822

Campbell CD, Ogburn EL, Lunetta KL, Lyon HN, Freedman ML, Groop LC, Altshuler D, Ardlie KG, Hirschhorn JN (2005) Demonstrating stratification in a European American population. Nat Genet 37: 868–72

Day FR, Loh PR, Scott RA, Ong KK, Perry JR (2016) A Robust Example of Collider Bias in a Genetic Association Study. Am J Hum Genet 98: 392–3

Devlin B, Bacanu SA, Roeder K (2004) Genomic Control to the extreme. Nat Genet 36: 1129–30; author reply 1131

Devlin B, Roeder K (1999) Genomic control for association studies. Biometrics 55: 997–1004

Franke A, Balschun T, Sina C, Ellinghaus D, Häsler R, Mayr G, Albrecht M, Wittig M, Buchert E, Nikolaus S, Gieger C, Wichmann HE, Sventoraityte J, Kupcinskas L, Onnie CM, Gazouli M, Anagnou NP, Strachan D, McArdle WL, Mathew CG, Rutgeerts P, Vermeire S, Vatn MH, Krawczak M, Rosenstiel P, Karlsen TH, Schreiber S (2010) Genome-wide association study for ulcerative colitis identifies risk loci at 7q22 and 22q13 (IL17REL). Nature Genetics 42: 292–294

Freedman ML, Reich D, Penney KL, McDonald GJ, Mignault AA, Patterson N, Gabriel SB, Topol EJ, Smoller JW, Pato CN, Pato MT, Petryshen TL, Kolonel LN, Lander ES, Sklar P, Henderson B, Hirschhorn JN, Altshuler D (2004) Assessing the impact of population stratification on genetic association studies. Nat Genet 36: 388–93

Guo J, Geng Z (1995) Collapsibility of Logistic Regression Coefficients. Journal of the Royal Statistical Society. Series B (Methodological) 1: 263–267

Helgason A, Yngvadottir B, Hrafnkelsson B, Gulcher J, Stefansson K (2005) An Icelandic example of the impact of population structure on association studies. Nat Genet 37: 90–5

Hirschhorn JN, Daly MJ (2005) Genome-wide association studies for common diseases and complex traits. Nat Rev Genet 6: 95–108

Hu P, Jiao R, Jin L, Xiong M (2018) Application of Causal Inference to Genomic Analysis: Advances in Methodology. Frontiers in Genetics 9

Klein RJ (2005) Complement Factor H Polymorphism in Age-Related Macular Degeneration. Science 308: 385–389

Lander ES, Schork NJ (1994) Genetic dissection of complex traits. Science 265: 2037–48

Leng S, Wu G, Collins LB, Thomas CL, Tellez CS, Jauregui AR, Picchi MA, Zhang X, Juri DE, Desai D, Amin SG, Crowell RE, Stidley CA, Liu Y, Swenberg JA, Lin Y, Wathelet MG, Gilliland FD, Belinsky SA (2015) Implication of a Chromosome 15q15.2 Locus in Regulating UBR1 and Predisposing Smokers to MGMT Methylation in Lung. Cancer Research 75: 3108–3117

Lohmueller KE, Pearce CL, Pike M, Lander ES, Hirschhorn JN (2003) Meta-analysis of genetic association studies supports a contribution of common variants to susceptibility to common disease. Nat Genet 33: 177–82

Marchini J, Cardon LR, Phillips MS, Donnelly P (2004) The effects of human population structure on large genetic association studies. Nat Genet 36: 512–7

Pearl J (2000) Causality: Models, Reasoning, and Inference. Cambridge University Press, Cambridge, UK

Petersen ML, van der Laan MJ (2014) Causal models and learning from data: integrating causal modeling and statistical estimation. Epidemiology 25: 418–26

Pirinen M, Donnelly P, Spencer CCA (2012) Including known covariates can reduce power to detect genetic effects in case-control studies. Nature Genetics 44: 848–851

Price AL, Patterson NJ, Plenge RM, Weinblatt ME, Shadick NA, Reich D (2006) Principal components analysis corrects for stratification in genome-wide association studies. Nat Genet 38: 904–9

Schisterman EF, Cole SR, Platt RW (2009) Overadjustment Bias and Unnecessary Adjustment in Epidemiologic Studies. Epidemiology 20: 488–495

Tam V, Patel N, Turcotte M, BosséY. ParéG, Meyre D (2019) Benefits and limitatio of genome-wide association studies. Nature reviews. Genetics

Thomas DC, Haile RW, Duggan D (2005) Recent developments in genomewide association scans: a workshop summary and review. Am J Hum Genet 77: 337–45

Tintle NL, Pottala JV, Lacey S, Ramachandran V, Westra J, Rogers A, Clark J, Olthoff B, Larson M, Harris W, Shearer GC (2015) A genome-wide association study of saturated, monoand polyunsaturated red blood cell fatty acids in the Framingham Heart Offspring Study. Prostaglandins, Leukotrienes and Essential Fatty Acids 94: 6572

Tomasetti C, Li L, Vogelstein B (2017) Stem cell divisions, somatic mutations, cancer etiology, and cancer prevention. Science 355: 1330–1334

Tomasetti C, Vogelstein B (2015) Variation in cancer risk among tissues can be explained by the number of stem cell divisions. Science 347: 78–81

Visscher PM, Brown MA, McCarthy MI, Yang J (2012) Five Years of GWAS Discovery. The American Journal of Human Genetics 90: 7–24

Zhang H, Chatterjee N, Rader D, Chen J (2018) Adjustment of nonconfounding covariates in case-control genetic association studies. Annals of Applied Statistics 12: 200–221

